# Effects of short-term salinity exposure on haemolymph osmolality, gill morphology and Na^+^/K^+^ - ATPase activity in Solenaia oleivora

**DOI:** 10.1101/2021.09.14.460389

**Authors:** Jingting Yao, Ting Zhang, Dongpo Xu, Guohua Lv, Wu Jin, Xueyan Ma, Yanfeng Zhou, Ruobo Gu, Haibo Wen

**Affiliations:** Freshwater Fisheries Research Centre, Chinese Academy of Fishery Sciences, Wuxi, China

**Keywords:** Solenaia oleivora, salinity exposure, ions concentration, gill morphology, Na^+^/K^+^-ATPase

## Abstract

In order to explore the physiological reaction to hyperosmotic environment, Solenaia oleivora were exposed to 2.23‰ salinity. In 48h, the hemolymph osmolality kept increasing, and the hemolymph protein concentration increased in the first 6h and then decreased significantly, while the free amino acid content increased in the first 24h and then kept stable (*P* < 0.05). The activity of Na^+^/K^+^-ATPase at 0h was significantly higher than other times in most organs except intestine, which was highest at 3h (*P* < 0.05). The ions concentration were also influenced. The concentration of Na^+^ rose in haemolymph, axe foot and intestine, but decreased in gill and hepatopancreas. In hemolymph, gill, hepatopancreases and adductor muscle, the K^+^ concentration was the highest at 0h, while in axe foot and intestine, it showed a positive tendency. The concentration of Cl^-^ in haemolymph, adductor muscle, intestine and axe foot were positively correlated with treatment time, while hepatopancreas showed opposite tendency. High salinity stress caused a difference in the gill histological structure, the gill structure shrunk, the gill lamellas space and shrinking degree showed an enlarging trend with salinity treatment time.

---

Solenaia belongs to the Mollusca phylum, Bivalvia, Palaeoheterodonta, Unionoida, Unionidae, which are endemic species in Asia. Nowadays, seven Solenaia species could be found in southeastern Asia, and four of them are recorded in China including S. carinatus, S. oleivorus, S. rivularis and S. triangularis (Heude, 1874-1885). Solenaia oleivora (Bivalvia, Unionidae, Gonideinae) is a proper rare freshwater bivalve distributed in the mid-lower Yangtze River and the middle Huai River drainage (Hu, 2005; Liu et al., 1979). It is an economically important freshwater mussel with high growth rate and high nutritional value (Xu et al., 2003; Yang et al., 2011). This species used to live in the relatively large ranges including Hebei province, Anhui province, Jiangsu province, Jiangxi province, Hubei province and Zhejiang province (Liu, 1979; Zhang and Qi, 1961). However, in recent years, the population size of S. oleivora declines dramatically for the reason of water pollution, habitat destruction, increasing capture pressure of the mussel and its host fishes, and only a few S. oleivora were observed in several lakes and rivers in Tianmen Hubei province, Poyang Lake Jiangxi province and Anhui province (Zhan et al., 2018; Li, 2012; Wen, 2009; Wang et al., 2015; Zhang et al., 2020; Xiong et al., 2012). As a result, the artificial propagation and artificial releasing have been closely concerned, nowadays, the difficulty of artificial propagation in S. oleivora was broken through, but the research on S. oleivora is limited (Zhang et al., 2020; Chen et al., 2020), and few data are available about its response to different environment (Xu et al., 2005).

Salinity is one of the dominant environmental factors influencing mussel physiological processes. Unfavourable salinity could influence several metabolic and physiological parameters in mussels including mortality (Bergman et al., 1996), growth rate (Wang and Qi, 2018), filtration rate (Hans et al., 2014), energy acquisition (Gardner and Thompson, 2001) and body health. The ability of existing at varying salinity depends on different species. Compared to marine mussels, freshwater mussels are basically able to live in water of a smaller range of salinity (Berger and Kharazova., 1997), and the process of salinity adaptations are less investigated.

Previous research reported that high salinity influences the energy metabolism of mussels by increasing oxygen consumption and ammonia excretion (Nie et al.,2017), promoting osmolarity and the concentration of plasma ions Na^+^, K^+^, Cl^-^ (Geng et al., 2016), as well as decreasing the activities of Na+/K+-ATPase (NKA) (Wang et al., 2019). NKA is a ubiquitous membrane-bound enzyme responsible for the transports of ions through cell membranes to regulate osmotic pressure and membrane permeability (Kulac et al., 2013). It is widely distributed in different tissues, driving the coupled extrusion of Na^+^ and uptake of K^+^ ions by hydrolyzing ATP. In high permeability medium, freshwater mussels are hypotonic to the medium, losing water and gaining salt passively. To compensate, they close their shells, decrease filtration rate and excrete more monovalent ions via the gills (Navarro., 1988; Cai et al., 2015).

According to our unpublished research, S. oleivora could tolerate the salinity of 2.23‰ without significant mortality. Therefore, a 48h-exposure trial to 2.23‰ saline water is designed to study the haemolymph osmolarity and the concentration of ions in different tissues to describe the response of S. oleivora to high permeability medium and assess the feasibility of salt-affected land utilization, and information on its response has implications on selecting the suitable sites for farming, artificial propagation and releasing.

## 2. Materials and methods

### 2.1 Experimental mussels and rearing conditions

One hundred and fifty (150) S. oleivora with an average initial weight of 4.84 ± 0.91 g were obtained from Freshwater Fisheries Research Center of the Chinese Academy of Fishery Sciences culture facility located at Nanquan, Wuxi, China.

Mussels were randomly distributed to 6 indoor tanks (25 specimens/tank) with volumes of 0.8 cubic meters each and 20-25 cm water depth. The mussels were fed with diluted Nannochloropsis (1mL/1L of water, Nanno 3600, Reed Mariculture Inc, Philippines) twice a day (8:30am and 16:30pm) for a one-week acclimatization period. Shell length (SL) and wet weight (WW) were determined before trial.

Water was changed twice a day after feeding for one hour and a half. Water quality and environmental indicators were closely monitored during the experiment. The tanks were supplied with fully aerated and filtered tap-water, and uninterrupted aerated to maintain an appropriate level of dissolved oxygen (> 5 mg/L), ammonium nitrogen (< 0.6 mg/L), and the water temperature varied 18-20°C. All procedures described in this study were in accordance with the policy of the Ethics Committee for Animal Experiments of the Nanjing Agricultural University.

### 2.2 Experimental method

The experiment was began after fasting for 24 hours. Two experimental salinities were designed, control group (freshwater) and salinity group (2.23‰, NaCl was diluted to the required salinity). Mussels in six tanks were conducted in triplicate with 25 mussels per replicate. Other environmental factors expect salinity were the same as acclimatization period.

### 2.3 Sample collection

The experiment was lasted for 48h, three mussels were sampled from each tank (9 specimens/group) randomly at 0, 3, 6, 12, 24, 48h. Haemolymph samples were obtained from the adductor muscle in the shell posterior, transferred to a 1.5 ml sterile micro-centrifuge tube, and stored at −20 °C for analysizing Na^+^/K^+^-ATPase activity, protein, chloride concentration, sodium concentration, potassium concentration and osmolality. Intestine, hepatopancreas, gill, adductor muscle and axe foot were then removed and washed with cold saline water (0.6 g/L), dried with filter paper, weighed and transferred to a 1.5 ml sterile micro-centrifuge tube, and storaged at −80 °C for RNA isolation, analyzing chloride concentration, sodium concentration, potassium concentration and Na^+^/K^+^-ATPase activity. For histology, one third of gills were fixed in Bouin’s solution for 24 h before embedding in paraffin according to standardized routine.

### 2.4 Na^+^/K^+^-ATPase activity, chloride, sodium, potassium in different tissues and haemolymph osmolality

Hepatopancreas, gills, adductor muscle, axe foot and intestinal samples were homogenized by adding sterile 0.6% saline solution to prepare 10% (W:V) homogenates, and then centrifuged at 4000×g for 10 min at 4°C. Supernate was used for further analyze in 12h.

NKA activity was analyzed according to a modified microtitre plate method from McCormick (1993), and microplate reader was used to measure the enzyme activity of NKA. Each sample was assayed in triplicate.

Na^+^, Cl^-^, and K^+^ concentrations in different tissues were measured using the commercially available kits (Nanjing Jiancheng Bioengineering Institute, Nanjing, China).

Haemolymph osmolality was measured by Osmomat 030 Cryoscopic Osmometer (Gonotec, Germany). For each analysis, 15μl of haemolymph was needed, and results were measured by comparing the temperature at crystallization point of the sample, pure water and standard solution.

Coomassie blue staining and calf plasma protein as a standard were used for haemolymph protein determination.

### 2.5 Slice preparation and intestinal index observation

Gills were fixed in Bouin’s solution for 24 h and embedded into paraffin blocks according to routine procedures, then cut into slices with a thickness of 7 μm, placed, stretched, dried and stained (Yao et al., 2019). The slices were then dehydrated with 95% and 100% ethanol followed by xylene and mounted with coverslips. Finally, the slices were observed via light microscopy to obtain the histological structure index.

### 2.6 Haemolymph free amino acids percentage

The free amino acids (FAAs) in haemolymph were quantified by the amino acid analyzer (Hitachi Model 835) according to the method of Lee et al. (2012).

### 2.7 Data analysis

Data were presented as means ± SD and analyzed via one-way analysis of variance (ANOVA) in SPSS version 17.0. Differences in mean values were analyzed via Duncan’s multiple range test when ANOVA identified differences among groups. The level of significance was set to *P* <0.05.

## 3. Results

### 3.1 Hemolymph osmolality and protein concentration

The data indicates that hemolymph osmolality and duration were positively correlated in 48h (*P* < 0.05, Figure 1). While the protein concentration increased significantly in the first 6h exposed to high salinity, and then decreased (*P* < 0.05, Figure 2).

**Figure 1.**
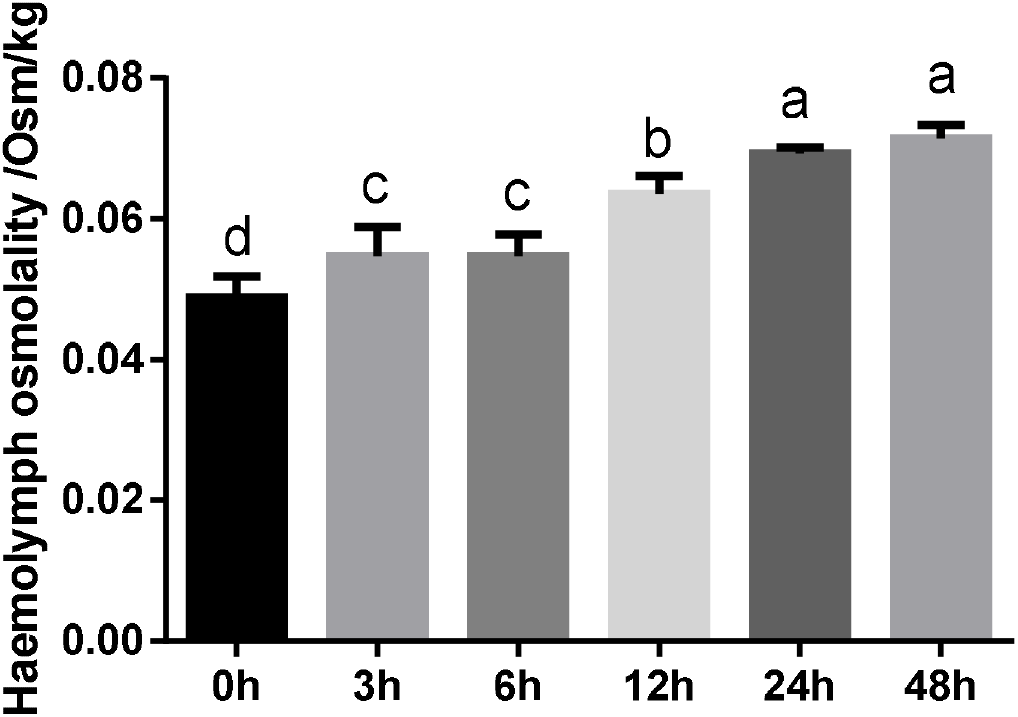
Hemolymph osmolality in S. oleivora reared at salinity Notes: Different letters indicate statistical differences (*P* < 0.05) between different time.

**Figure 2.**
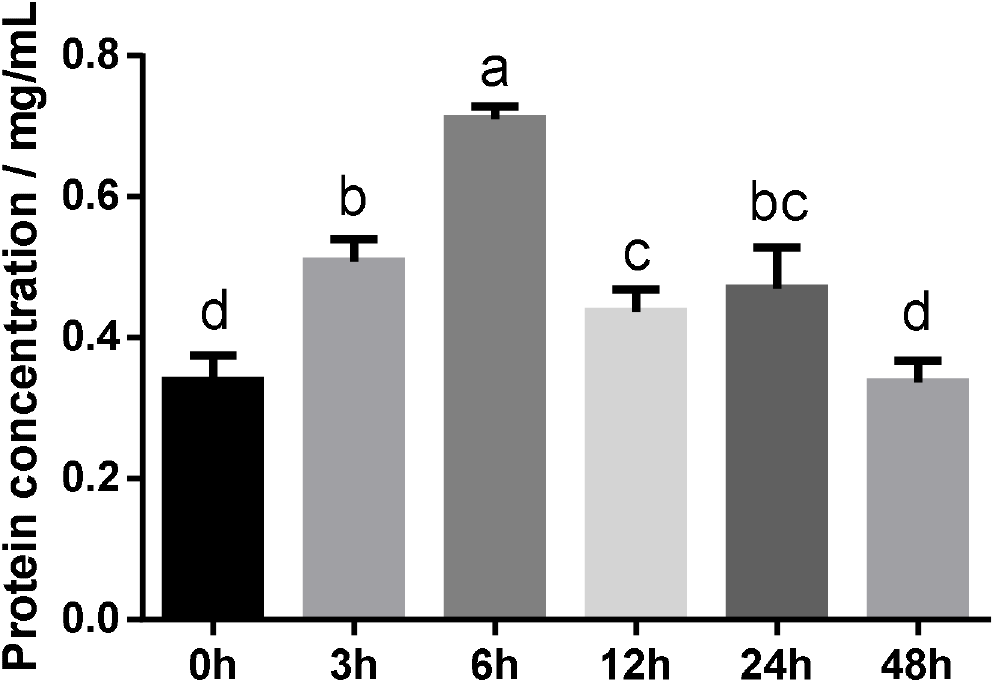
Hemolymph protein concentration in S. oleivora reared at salinity Notes: Different letters indicate statistical differences (*P* < 0.05) between different time.

### 3.2 Na^+^/ K^+^-ATPase activity

In gill, the activity of NKA at 0h was significantly higher than other times (*P* < 0.05, Figure 3). In adductor muscle, hepatopancreas and axe foot, the activity of NKA decreased significantly with the time exposed to salinity (*P* < 0.05, Figure 3). While in intestine, the activity of NKA increased and then decreased at 12h (*P* < 0.05, Figure 3).

**Figure 3.**
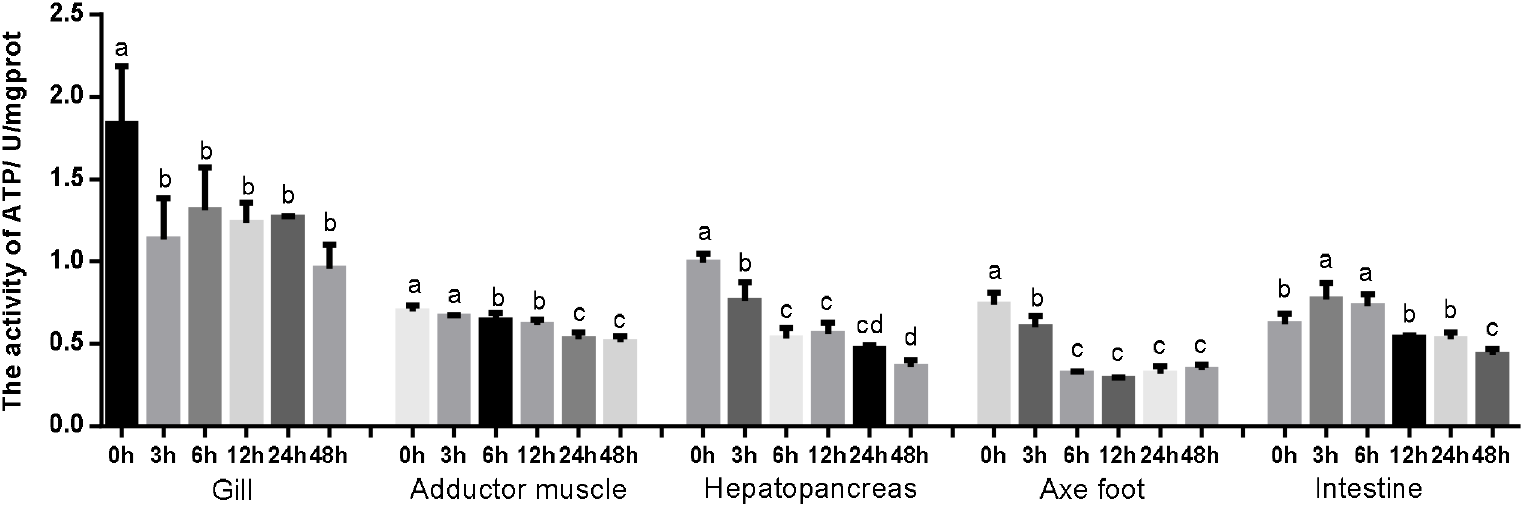
Na^+^/ K^+^-ATPase activity (U/mgprot) in different tissues of S. oleivora reared at salinity Notes: Different letters in the same tissue indicate statistical differences (*P* < 0.05) between different time.

### 3.3 The concentration of Na^+^, K^+^, Cl^-^ in different tissues

In hemolymph, the concentration of Na^+^ at 24h and 48h were significantly higher than before (*P* < 0.05, Figure 4). In gill and hepatopancreas, the concentration of Na^+^decreased in the first 6h and then kept stable (*P* < 0.05, Figure 4). In adductor muscle, the concentration of Na^+^ increased at 12h and 24h and then decreased (*P* < 0.05, Figure 4). In axe foot and intestine, the Na^+^ concentration at 48h was significantly higher than that before (*P* < 0.05, Figure 4).

**Figure 4.**
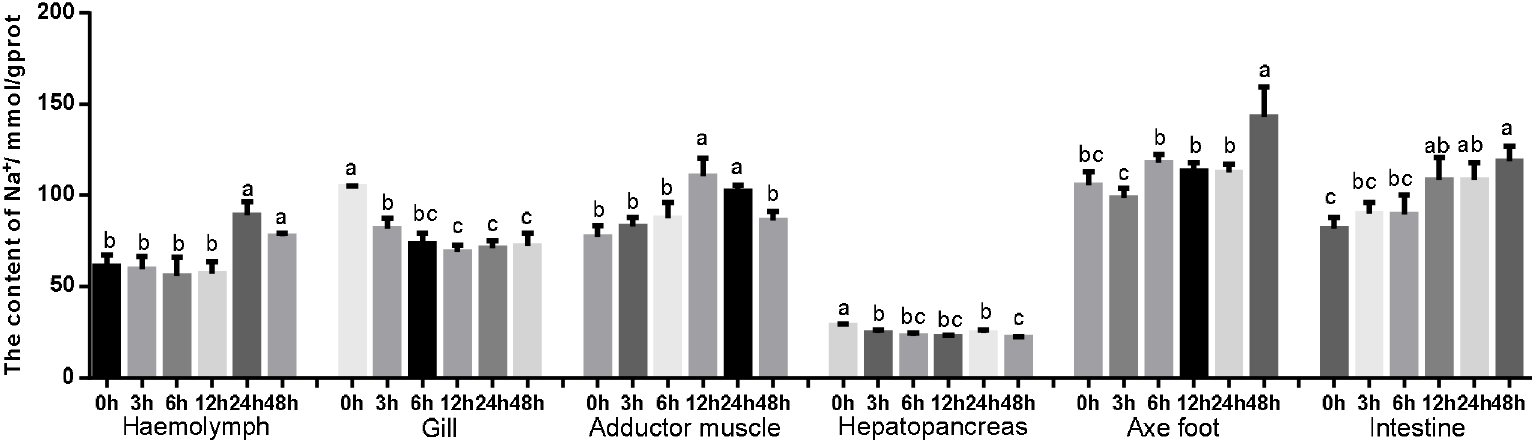
Concentrations of Na^+^ (mmol/gprot) in different tissues of S. oleivora reared at salinity Notes: Different letters in the same tissue indicate statistical differences (*P* < 0.05) between different time.

In hemolymph, gill and adductor muscle, the concentration of K^+^ at 0h was significantly higher than that later (*P* < 0.05, Figure 5), and K^+^ in hepatopancreases at 0h was significantly higher than that at 48h (*P* < 0.05, Figure 5). While in axe foot, the K^+^ concentration increased in 48h, and in intestine, it increased significantly in the first 6h and then kept stable (*P* < 0.05, Figure 5).

**Figure 5.**
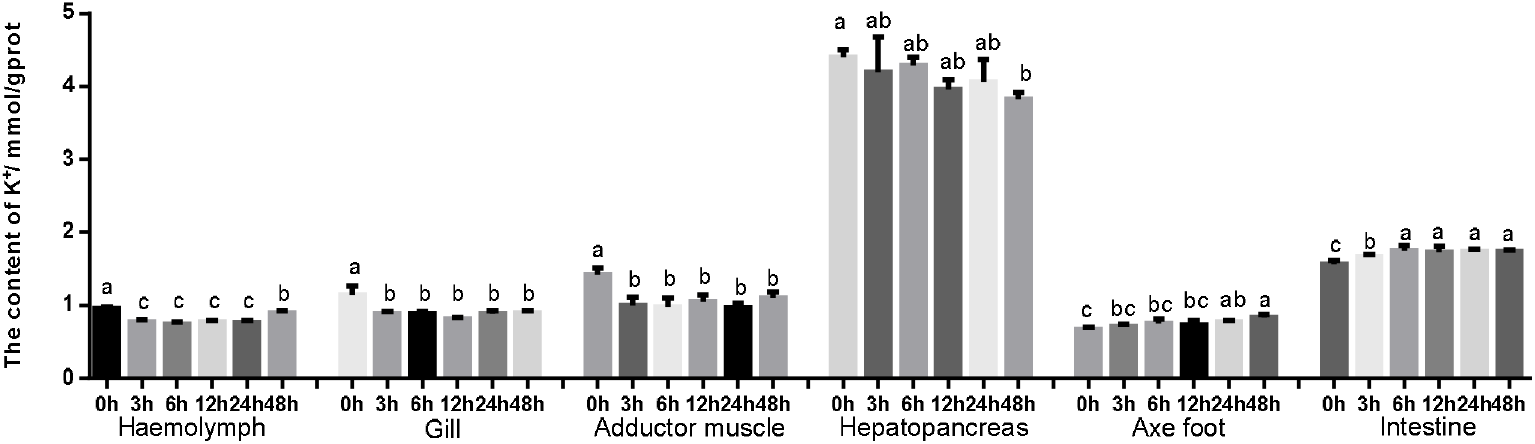
Concentrations of K^+^ (mmol/gprot) in different tissues of S. oleivora reared at salinity Notes: Different letters in the same tissue indicate statistical differences (*P* < 0.05) between different time.

In hemolymph, the concentration of Cl^-^ kept stable in the first 6h and then increased significantly (*P* < 0.05, Figure 6). In gill, it increased at 6h and decreased at 48h (*P* < 0.05, Figure 6). In adductor muscle and intestine, the concentration of Cl^-^ at 0h was significantly lower than that later (*P* < 0.05, Figure 6). In axe foot, the Cl^-^ concentration kept stable in the first 3h and then increased, while in hepatopancreas, it significantly decreased after 24h (*P* < 0.05, Figure 6).

**Figure 6.**
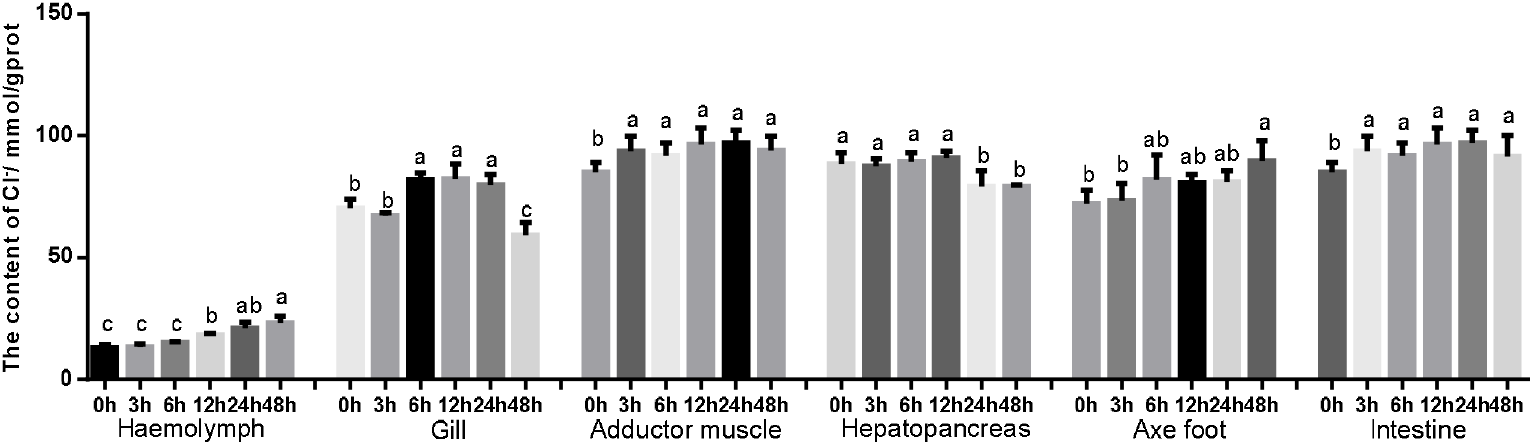
Concentrations of Cl^-^ (mmol/gprot) in different tissues of S. oleivora reared at salinity Notes: Different letters in the same tissue indicate statistical differences (*P* < 0.05) between different time.

### 3.4 Gill histomorphology

The results showed that high salinity stress induced alterations of gill histomorphology (Figure 7). The gill filaments of S. oleivora were closely arranged at the beginning, the top of the gills expanded into a rod shape. After 3h high salinity treatment, the gill structure shrunk, the gill lamellas space increased and the breadth of the gill filaments decreased. Six hours later, the intervals of the gill lamella and shrinking degree showed an enlarging trend with the increase of stress hours.

**Figure 7.**
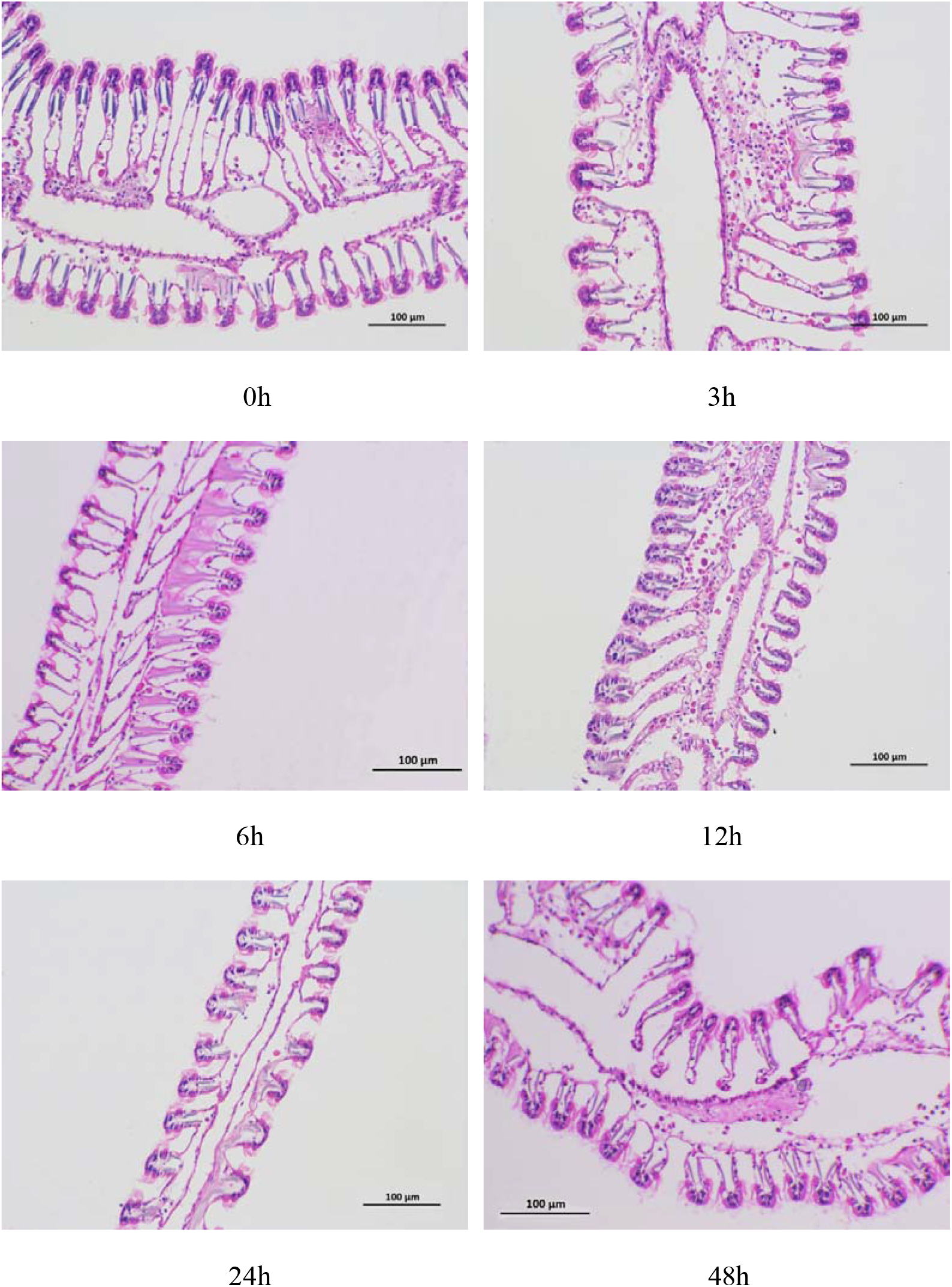
the structure of gill and chloride cell

### 3.5 Hemolymph free amino acids content

Significant (*P* < 0.05) difference was observed in the content of essential amino acids (EAA), non-essential amino acids (NEAA) and total amino acids (TAA) in hemolymph, except Cys, Ile, Phe and His (Table 1). The content of NEAA and TAA increased significantly in the first 24h and then kept stable (*P* < 0.05), while the EAA increased in the first 12h, decreased at 24h and increased again at 48h (*P* < 0.05).

**Table 1.**
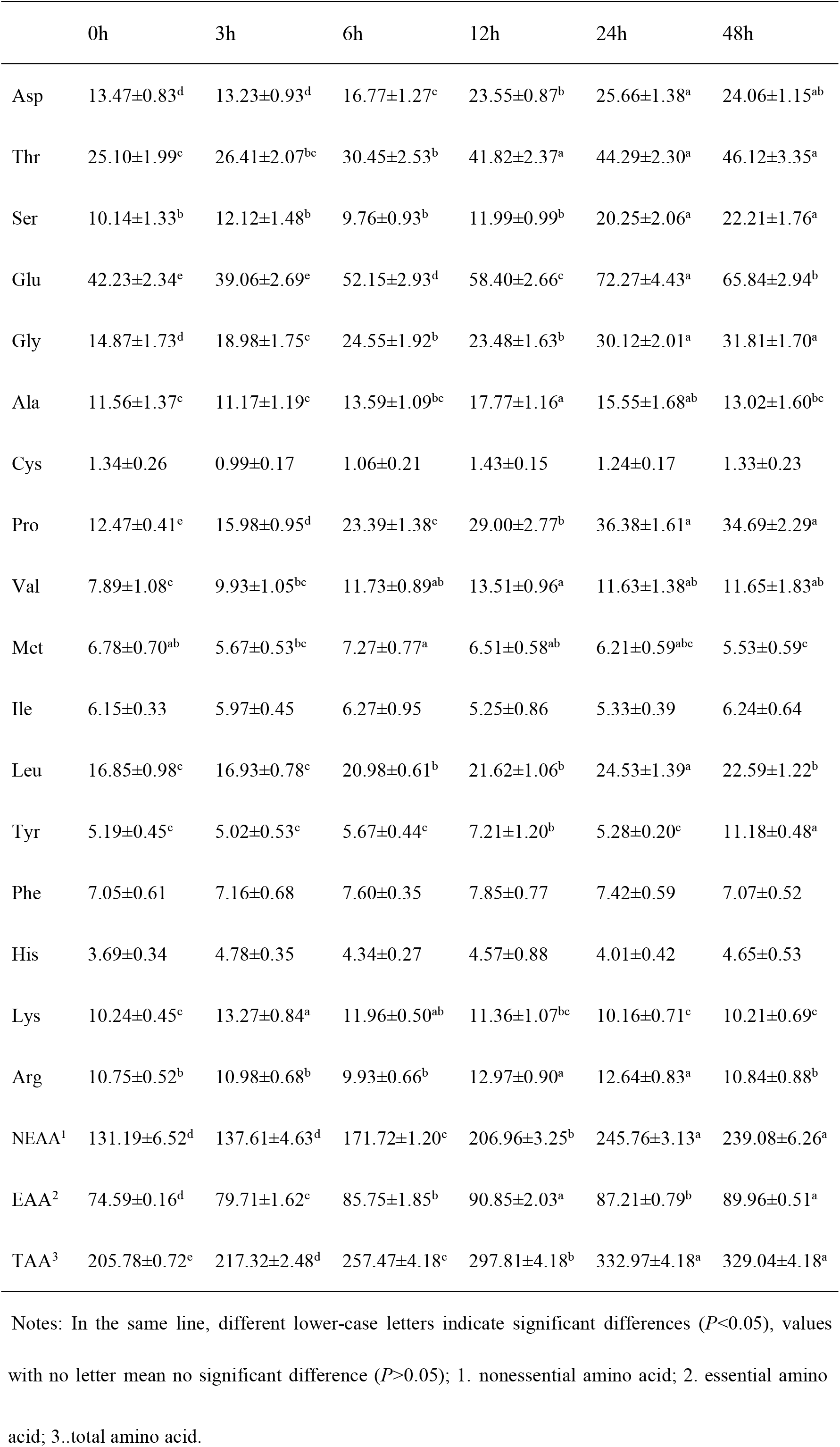
Free amino acid content in hemolymph (mg/L) of S. oleivora (means ± SD, n= 3)

## 4. Discussion

Salinity is a crucial environmental factor influencing mussels’ physiological processes including osmotic pressure, antioxidant ability, feeding activity, etc (Navarro, 1988; Shumway, 1977). Most aquatic invertebrates such as S. oleivora are not equipped with advanced mechanisms to regulate their internal environment, therefore, osmotic adaptation plays an essential role (Hosoi, 2005). Typically, when a mussel enters high salinity water, firstly, it maintains the osmotic pressure at a normal level by closing the shell and blocking the water-salt exchange with the environment, and then, the osmotic pressure increases for the reason of passive water loss, finally, the ion discharge mechanism is activated, which regulates osmotic pressure to be stable or level off gradually (Berger and Kharazova, 1997). Cai et. al (2015) proved that under abrupt stress at salinity of 40‰, the hemolymph osmolality of juvenile ark shell (*Anadara broughtonii*) increased significantly in 48h and then kept stable.

Most aquatic invertebrates maintain free amino acid pools to adapt to external osmotic changes, and several researches have indicated the relationship between osmotic stress and free amino acid pool in aquatic species (Zurburg and Zwaan, 1981; Fu et al., 2017). In aquatic species, FAA and protein are important end products, substrates or intermediates of energy metabolism (Berger and Kharazova, 1997). Previous study indicated that the metabolism of protein in aquatic invertebrates decreases under hyperosmotic environment, while hypoosmotic stress could promote the biosynthesis and accumulation of amino acids (Navarro, 1988; Matsushima et al., 1987). In the present study, the content of hemolymph protein increased and then decreased at 12h, and most of hemolymph FAA increased significantly in 48h after treated by high salinity, such as aspartic acid, serine, threonine, glutamic acid. Similar results were presented in other molluscs. Yang et al. (2016) reported that the protein catabolism of *Meretrix meretrix* increased when external salinity was higher than 20‰, while the metabolism of carbohydrate and fat receded. Hosoi et al. (2003) indicated that under hyperosmotic stress, the free amino acids content of glycine, alanine, b-alanine, proline, arginine and taurine in the mantle of the Pacific oyster (*Crassostrea gigas*) increased significantly 72h (Hosoi et al., 2003). The improvement of free amino acids concentration may result from high proteolytic activities, which hydrolyze protein to regulate hemolymph osmatic pressure, as well as provide energy to adapt to environment (Siebers et al., 1972; Farmer and Reeve, 1978). Therefore, we suppose that under hyperosmotic environment, S. oleivora promotes proteolysis activity to improve FAA content, and the hemolymph FAA content plays important role in high salinity adaptations.

The main factors regulating osmotic pressure are ions concentrations and bioorganic compounds, such as protein, amino acids and other organic molecules. Na^+^and Cl^-^ are the crucial osmosis-related ions in hemolymph, whose contributions are more important than proteins and FAA. Cheng et al. (2002) reported that in Taiwan abalone *Haliotis diversicolor supertexta*, the concentration of hemolymph Na^+^, K^+^ and Cl^-^ showed positive correlation with hemolymph osmolality, and Na^+^ and Cl^-^ contribute to 47.7–51.6% and 37.1–42.2% of the hemolymph osmotic concentration respectively. Natochin et al. (1979) believed that at high salinities, the passive influx of Na^+^ and Cl^-^ is automatic in mollusc, which is a defense reaction to protect the stabilization of cell volume. Similar results were obtained in the present study, Na^+^ and Cl^-^ concentrations were significantly increased in hemolymph, axe foot, adductor muscle and intestine, while in hepatopancreas, Na^+^ and Cl^-^ concentrations decrease with time goes by. The special functions of different tissues might be an important reason. Hepatopancreas is the metabolism center in mollusc, under external stress, the synthesis of related enzymes and other macromolecule would be activated, and decomposition of nutrients would be improved to resist stress such as unproper osmolality and anoxia, and it may be connected with the excretion of ions (Lama et al., 2013; Silva-Pando et al., 1978).

NKA is a trans-membrane protein distributed in diverse tissues and cells in most animals. It uses the energy derived from ATP hydrolysis, driving the extrusion of Na^+^ and uptake of K^+^ ions against their electrochemical gradients to maintain the membrane potential. In aquatic species, a crucial function of NKA is to regulate osmotic pressure. Numerous researches reported that NKA is responsible for the plasma osmolality stability at high-salinity levels (Arjona et al., 2007; Palacios et al., 2004; Zhang et al., 2017). Wang and Cao (2019) indicated that in 48h, the activity of visceral NKA in Scallop (*Crassadoma gigantea*) under high salinity was lower than normal level especially in the case of sudden salinity change. Similar result was also reported in *Cyclina sinensis* and ark shell (*Anadara broughtonii*), the activity of NKA in gill decreased in 14d and 168h respectively (Cai et al., 2015; Lin et al., 2012). Qian et al. (2015) believe that the activity of NKA keeps the highest level at optimal external salinity. In the present study, the activity of NKA decreased in most tissues expect intestine, which increased in the first 6h and then decreased. A great number of researches believe that increased NKA activity promotes the excretion of excess salt ions to maintain normal ion concentration (Liu et al., 2008; Zhao et al., 2006). But salinity is not the only factor affecting NKA activity, different regulatory mechanism may be related in different species (Xu et al., 2007). Geng et al. (2016) indicated that the aquatic species improve NKA activity, discharge excess salt ions from the body when there was a large difference between plasma osmotic pressure in vivo and water osmotic pressure in vitro, and the NKA activity would become smaller when the difference decreased. Our previous study indicated that the osmolality of S. oleivora is 6‰, which is higher than freshwater. Therefore, we suspect that the gill NKA activity of S. oleivora decreased when transported from freshwater to 2.23‰ because less ions are needed to be absorbed. Apart from that, as the hemolymph Na^+^ and Cl^-^ concentrations increased significantly, the NKA activity in internal organs decreased, slow down the excretion of Na^+^ from tissues to hemolymph to regulate osmolality. In addition, Yu et al. (2014) believed that improper external osmolality damaged cell membrane, induced apoptosis, it could be a reasonable cause for reduced NKA activity as well.

Intestine is a key organ absorbing nutrition, water, and ions in aquatic species (Usher et al., 1991). The absorption and transport of NaCl against the gradient of concentrations fluid is accomplished essentially by the electrogenic basolateral NKA (Tresguerres et al., 2010). Previous study has demonstrated that the intestinal NKA activity in aquatic species enhanced at higher external salinities (Ruiz-Jarabo et al., 2017). It is generally accepted that the ingestion of salt ions is restrained and the absorption of water is improved under high salinity stress. However, Li et al. (2014) believed that *Oreochromis mossambicus* could increase the expression of NKA in intestinal tract to response to increasing salinity challenge, and it may be related to the facilitation of water uptake (Li et al., 2014). This may partly explain the improvement of NAK activity in intestine in the first 6h. Furthermore, as S. oleivora needs more energy to regulate osmolality, more nutrition may be absorbed thought intestine, and the activity of NKA might be facilitated to provide more energy (Hu et al., 2019; Li et al,. 2019).

Gill is the major organ responsible for osmoregulation in aquatic species. The gill epithelia contain high level of NKA, which provides the energy for many transporting systems to maintain intracellular homeostasis (Marshall, 2002). In freshwater bivalve, which are hypertonic to the environment, the diffusional loss of ions and the osmotic influx of water are balanced by gills’ ions absorption. In the present study, however, in hypertonic medium, lower requirement for ion pumping reduced the activity of ATP, that may be the crucial reason for the reduction of NKA in gill under salinity stress. The variation of gill histomorphology also proved it, the gill lamellae became narrowed and shorter, and the surface area for ions-exchange decreased as well.

In summary, the present study described the osmolality-related physiological reaction of S. oleivora under high environmental salinity. From the data of our experiment, the osmolality of S. oleivora increased and kept stable in 48h. This will be provided a basis for the artificial propagation, artificial releasing and initial adaptation of S. oleivora. Further research is needed to determine whether salinity has detrimental influence on the growth performance and health of S. oleivora, and other environment factors should be concerned as well.

